# The value of genotype-specific reference for transcriptome analyses

**DOI:** 10.1101/2021.09.14.460213

**Authors:** Wenbin Guo, Max Coulter, Robbie Waugh, Runxuan Zhang

**Affiliations:** Information and Computational Sciences, James Hutton Institute, Dundee DD2 5DA, Scotland, UK; Plant Sciences Division, School of Life Sciences, University of Dundee at The James Hutton Institute, Invergowrie, Dundee DD2 5DA, Scotland, UK; Cell and Molecular Sciences, James Hutton Institute, Dundee DD2 5DA, Scotland, UK

## Abstract

High quality transcriptome assembly using short reads from RNA-seq data still heavily relies upon reference-based approaches, of which the primary step is to align RNA-seq reads to a single reference genome of haploid sequence. However, it is increasingly apparent that while different genotypes within a species share “core” genes, they also contain variable numbers of “specific” genes that are only present a subset of individuals. Using a common reference may thus lead to a loss of genotype-specific information in the assembled transcript dataset and the generation of erroneous, incomplete or misleading transcriptomics analysis results. With the recent development of pan-genome information in many species, it is important that we understand the limitations of single genotype references for transcriptomics analysis. In this study, we quantitively evaluated the advantages of using genotype-specific reference genomes for transcriptome assembly and analysis using cultivated barley as a model. We mapped barley cultivar Barke RNA-seq reads to the Barke genome and to the cultivar Morex genome (common barley genome reference) to construct a genotype specific Reference Transcript Dataset (sRTD) and a common Reference Transcript Datasets (cRTD), respectively. We compared the two RTDs according to their transcript diversity, transcript sequence and structure similarity and the accuracy they provided for transcript quantification and differential expression analysis. Our evaluation shows that the sRTD has a significantly higher diversity of transcripts and alternative splicing events. Despite using a high-quality reference genome for assembly of the cRTD, we miss ca. 40% transcripts present in the sRTD and cRTD only has ca. 70% true assemblies. We found that the sRTD is more accurate for transcript quantification as well as differential expression and differential alternative splicing analysis. However, gene level quantification and comparative expression analysis are less affected by the source RTD, which indicates that analysing transcriptomic data at the gene level may be a reasonable compromise when a high-quality genotype-specific reference is not available.

## Introduction

For more than a decade, RNA sequencing (RNA-seq) has become the preferred method for large scale transcript identification and quantification (Wang *et al*., 2009; Conesa *et al*., 2016). It accesses a more diverse collection of transcripts than earlier technologies like microarrays and allows studies of post transcriptional regulation, for example by alternative splicing (AS) (Mantione *et al*., 2014; Conesa *et al*., 2016; Zhao, 2019). Consequently, one of the main uses of RNA-seq data is to quantify gene expression at both gene and transcript levels. Several studies have shown that for gene level analysis, quantification at transcript resolution improves the overall estimation of gene expression (Zhao *et al*., 2015; Trapnell *et al*., 2013). Currently, the gold standard quantification of transcripts depends upon the use of well-annotated reference transcript datasets (RTDs) (Zhang *et al*., 2017; Brown *et al*., 2017; Zhang *et al*., 2015; Rapazote-Flores *et al*., 2019) in conjunction with rapid and accurate computational programs that implement approaches based on *pseudoalignment*, such as Salmon (Patro *et al*., 2017) and Kallisto (Bray *et al*., 2016). *De novo* assembly methods can assemble transcripts without guidance of a reference genome, but they suffer from significantly enhanced mis-assemblies and low sensitivities (Martin and Wang, 2011; Marchant *et al*., 2016; Conesa *et al*., 2016). High quality transcript assembly is still frequently derived from genome reference mapping-based assembly approaches.

In the genome reference mapping-based approach, transcript assembly generally starts by mapping RNA-seq reads to a common reference; a haploid sequence considered representative of the genomes of related individuals within a phylogenetic clade (in our example this would be all domesticated barley genotypes). For example the latest barley RTD BaRTv1.0 was constructed by mapping RNA-seq reads from over 150 barley cultivars to the reference Morex genome (Hv_IBSC_PGSB_v2) (Rapazote-Flores *et al*., 2019). However pan-genome studies have revealed that diverse genotypes contain a shared set of core genes as well as a large proportion (20 – 50%) of genotype-specific genes in subsets of individuals (Li *et al*., 2014; Golicz *et al*., 2016; Montenegro *et al*., 2017; Hirsch *et al*., 2014; Jin *et al*., 2016; Sun *et al*., 2017). This variation is even more complex at the transcript level when considering post transcriptional modifications such as alternative splicing (AS). Moreover, genomic sequences of common reference and individual strains contain frequent sequence variations, such as single nucleotide polymorphisms (SNP), short deletions and insertions (INDEL), which also affect the transcript determinations in the assembly (Munger *et al*., 2014); sequence variation at splice sites will disrupt the recognition of introns and exons and alter protein translations (Anna and Monika, 2018; Baeza-Centurion *et al*., 2020).

Recently, reference quality sequence assemblies for 20 genotypes of diverse geographical origin, spike morphology and annual growth habit have been made available through investigations into variation in the barley pan-genome (Jayakodi *et al*., 2020). In this study, ∼1.5 million of Present/absent variants (PAVs) ranging from 50bp to ∼1Mbp were identified and 5,602 deletions longer than 5 kilobases (kb) were found in Barke relative to Morex alone. Thus, the impact of moving from common reference to genotype specific genome reference for transcriptomics studies, such as transcript assembly, quantification and differential expression analysis could be profound. The availability of high-quality genotype specific genome references has created an opportunity to investigate the impacts genotype specific genetic variation on transcriptomics studies.

Here we present a comprehensive investigation to quantitatively explore the benefits of using a genotype-specific reference genome for transcriptomics analysis in barley. We mapped RNA-seq data generated from Barke to the reference Morex genome (common reference) as well as a newly assembled high quality Barke genome (genotype-specific genome) and generated a common reference based RTD (cRTD) and genotype-specific RTD (sRTD) in parallel using the same tools and parameters (Fig 1A). We evaluated the impact of using the sRTD in comparison to cRTD on transcriptome analysis using the following metrics: 1) gene and transcript diversity; 2) transcript sequences; 3) transcript structures; 4) alternative splicing; 5) quantification of transcript abundance and 6) differential expression accuracy (Figure 1B).

**Figure 1:**
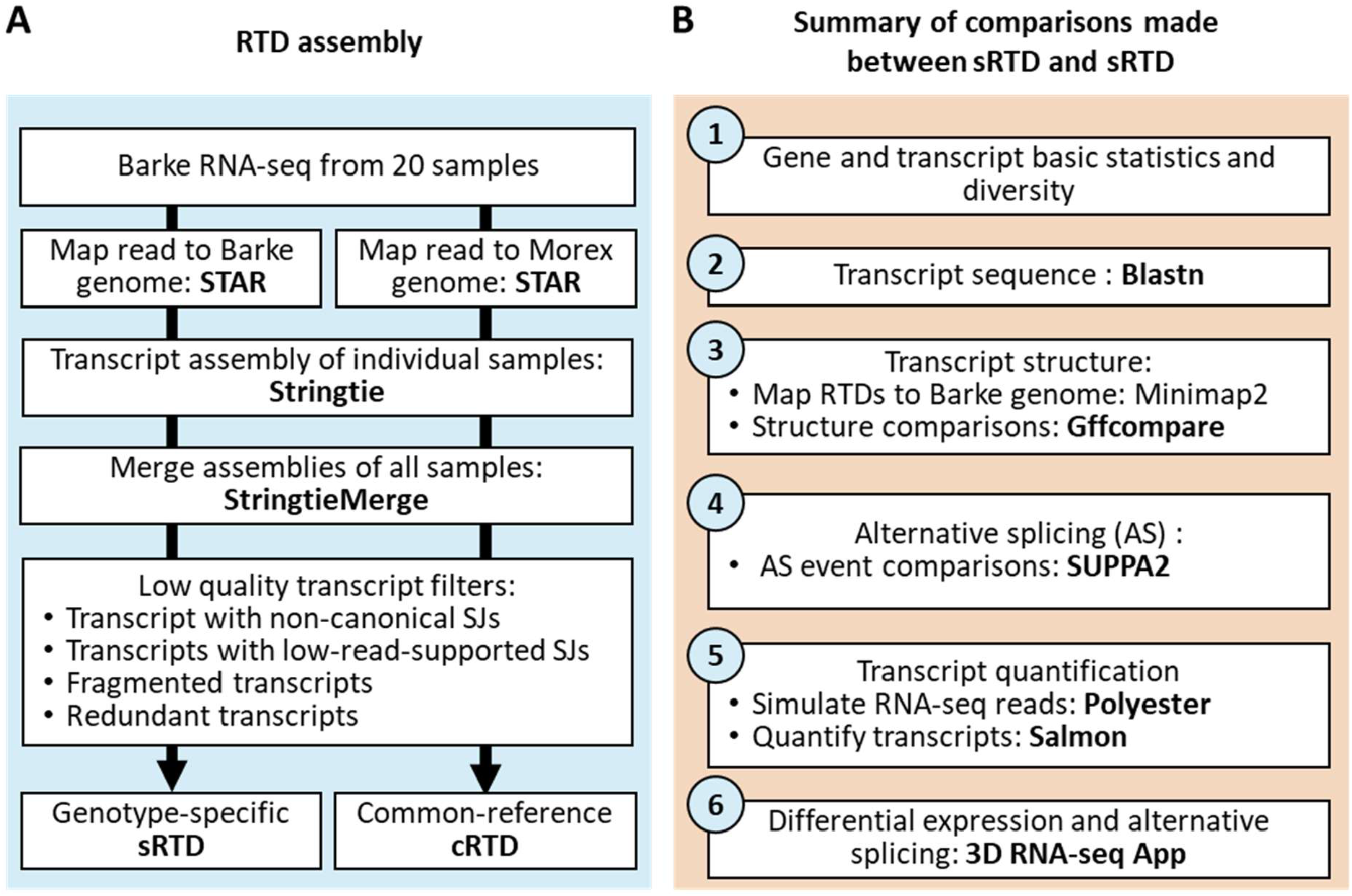
Workflow of genotype specific RTD studies. (A) RTD assembly pipeline for Barke RNA-seq read mapping to Barke and Morex genome, respectively. (B) Evaluation pipeline of the assembled sRTD and cRTD.

## Methods

### Data pre-processing and transcriptome assembly

Barke Illumina RNA-seq data from 20 tissues were taken from Coulter *et al*., (2021). The adapters on the raw RNA-seq reads were trimmed using Trimmomatic v0.39 (Bolger *et al*., 2014) with quality parameters chosen with the guidance of FastQC v0.11.9 (http://www.bioinformatics.babraham.ac.uk/projects/fastqc/). The trimmed reads in each sample were mapped to the Barke genome (Jayakodi *et al*., 2020) and Morex genome (Mascher *et al*., 2017) with STAR v2.7.3a (Dobin *et al*., 2013). We implemented the two-pass approach to increase the sensitivity of splice junction (SJ) discovery. The minimum and maximum intron sizes were set to 60 and 15,000 in both passes. We allowed 2 mismatches at the first pass to improve the SJ detection sensitivity. The detected SJs in the first pass mapping were merged across samples and used as guidance for read alignment in the second pass. To reduce spurious alignments caused by sequence variation and compare the RTDs more precisely, mismatch was not allowed in the second pass of read mapping. We assembled the sRTD and cRTD in parallel using the same processes and parameters using Stringtie v2.1.4 (Pertea *et al*., 2015). The assemblies of 20 samples were merged with Stringtie-merge (Pertea *et al*., 2015). We applied various filters to achieve high quality assemblies. 1) The transcripts with either non-canonical SJs or low read support (support criteria ≥ 3 uniquely aligned reads in ≥ 2 samples) were filtered out. 2) Redundant transcripts were defined as those with the same intron combinations but with different first and/or last exon lengths. Only the longest transcript amongst the same group of redundant transcripts was kept. 3) Transcript fragments < 70% of the length of the longest in the same group of redundant transcripts were removed. To determine protein-coding transcripts, we extracted the transcript sequences of sRTD and cRTD from the Barke and Morex genomes with Gffread (Pertea and Pertea, 2020). We used Transuite, which identified authentic AUGs in transcript sequences, to translate them into protein sequences (Entizne *et al*., 2020). We queried the protein sequences against the plant protein database UniProtKB for best-hit of annotated proteins by using Blastp (Schneider *et al*., 2009). To identify alternative splicing (AS), we employed SUPPA2 to demonstrate the diversity of AS events of retained intron (RI), alternative 5’ splice-site (A5), alternative 3’ splice-site (A3), skipping exon (SE), alternative first exon (AF), alternative last exon (AL) and mutually exclusive exons (MX) (Trincado *et al*., 2018).

### Assessing the impact on transcript sequences and structures

We aligned the transcript sequences of both cRTD and sRTD (query) to transcript sequences of sRTD (target) with Blastn. Blasting sRTD against sRTD was used as a reference to estimate the frequency that Blastn fails to align a query sequence correctly. Each query returned only one hit with the best e-value and bit-score (e-value must < 1.0e-5 and Blastn parameter max_target_seqs = 1). If a target was aligned by multiple query transcripts, only the one/ones with the maximum matched bases were retrieved. To further evaluate the proportions of matches to the target and query lengths, we treated the query-to-target matched bases as true positives (TP) of sequence alignment. The unaligned bases of the target and query were false negatives (FN) and false positives (FP), respectively. The proportions of sequence overlap were evaluated with precision (TP/(TP+FP)) and recall (TP/(TP+FN)), and their weighted mean F1 score (2×(Recall × Precision) / (Recall + Precision)) which took both recall and precision into account.

As sRTD and cRTD were assembled from read mappings to different genome references, their coordinates and structures cannot be compared readily. In order to compare the structures of transcripts at equivalent genome coordinates, we aligned the transcript sequences of both cRTD and sRTD to the Barke genome with Minimap2 v2.17 (Li, 2018), denoting the results as cRTD.minimap2 and sRTD.minimap2. Mis-aligned transcripts were filtered if the fragments of a transcript aligned to multiple chromosomes or strands and the aligned transcripts did not meet the criteria of minimum intron size 60 and maximum intron size 15,000. Transcript structures were evaluated by querying the genome coordinates of cRTD.minimap2 and sRTD.minimap2 against target sRTD at levels of nucleotide bases, introns, intron chains (combination of introns), exons, transcripts and gene loci by using GffCompare v0.11.2 (Pertea and Pertea, 2020). SUPPA2 (Trincado *et al*., 2018) was also used to investigate AS event and percent spliced-ins (PSIs) in annotations of cRTD.minimap2 and sRTD.minimap2. The AS events and PSIs of these two annotations were compared to that of sRTD. At the AS event level, we defined True Positives (TP) as the shared events between query (cRTD.minimap2 or sRTD.minimap2) and target (sRTD), False Positives (FP) as the events only in the query and False Negatives (FN) as the events only in target. Then, precision and recall were used to measure agreement between query and target. The AS event PSI values were calculated based on the transcript per million reads (TPMs) from the Salmon quantification of sRTD.minimap2 and cRTD.minimap2 with Barke RNA-seq reads from all 20 samples. In these analyses, the comparison of sRTD.minimap2 to sRTD provided a reference to estimate technical errors in Minimap2, GffCompare and SUPPA2.

### Assessing the impact on the accuracy of transcript quantification

To quantify the absolute quantification accuracy using both RTDs, we also used simulated RNA-seq reads to compare the quantifications between sRTD and cRTD at both transcript and gene levels. Specifically, the numbers of simulated reads for the sRTD transcripts were directly specified according to a read count matrix. To have replications and make the simulation results reflect real experimental data as much as possible, we used the read count matrix from the transcript quantification of “caryopsis” and “root” tissues, which was a subset of the Barke RNA-seq seven-tissue data in Jayakodi et al., (2020), each with 3 biological replicates. We used the Polyester R package to simulate 150-bp paired-end reads from the sRTD transcript sequences according to the read count matrix (Frazee *et al*., 2015). Then we quantified the cRTD and sRTD with Salmon by using the simulated RNA-seq reads (Patro *et al*., 2017). The gene and transcript quantifications of two RTDs were compared, in terms of Pearson and Spearman correlation and relative errors, to the ground truth read count matrix used to generate the data.

### Assessing the impact on differential gene and alternative splicing analysis

We applied the 3D RNA-seq pipeline (Guo *et al*., 2019) to analyse the differentially expressed (DE) genes and transcripts, differential alternative splicing (DAS) and differential transcript usage (DTU) based on three datasets. These were sRTD and cRTD quantifications from the simulated RNA-seq reads and the read count matrix, which is the ground truth for the simulation. In all analyses, we set the contrast group for testing expression changes to “caryopsis vs root”. The significance was determined with a Benjamini-Hochberg (BH) adjusted p-value <0.01 and absolute log2-fold-change ≥ 1 for DE genes and transcripts and BH adjusted p-value <0.01 and absolute Delta present spliced (Δ*PS*) ≥ 10% for the DAS genes and DTU transcripts. To compare the quantification at individual gene and transcript level, we used the results of transcript sequence and structure comparisons to match the equivalent individuals.

## Results

### Genome mapping comparisons

We used STAR to map Barke RNA-seq reads to the Barke and Morex genomes. The average mapping statistics of the samples are shown in Table 1. We can see that > 8.6 million extra pairs of reads (7.26% more of total) are uniquely mapped to the Barke genome compared to Morex. The percentage of unmapped reads is higher when mapping to the common reference (Morex). The majority of the unmapped reads (> 98%) are due to the mapping length being less than 2/3 of the mapped read length. STAR statistics show that 14.13% of total reads are unmapped in the common reference for this reason, while this number dropped to 6.90% in the genotype specific (Barke) genome. The genotype specific genome reference has less of these missing segments than common reference thus having a low percentage of unmapped reads. The spliced reads used to locate introns in the transcript assembly are also more abundant – by >9.4 million pairs of reads (10.96%) -in the genotype specific alignment. Sequence variations including deletions and insertions, increase from 0.01% to 0.02% and 0.00% to 0.02% respectively with their average lengths longer when mapping to the common reference (Table 1)

**Table 1:** Average statistics of STAR read mapping to Barke and Morex genomes.

|  | Category | Barke (genotype-specific ref) | Morex (Common ref) |
| --- | --- | --- | --- |
| Mapping | Uniquely mapped reads number | 107,770,451.15 (89.89%) | 99,142,457.6 (82.63%) |
|  | Average mapped length | 288.64 | 279.57 |
|  | Number of reads mapped to multiple loci (<10 loci) | 3,271,287.2 (2.64%) | 3,656,786.55 (3.01%) |
|  | Number of reads mapped to too many loci (<10 loci) | 534,171.1 (0.46%) | 131,798.65 (0.12%) |
|  | Number of reads unmapped: too many mismatches | 0 (0.00%) | 0 (0.00%) |
|  | Number of reads unmapped: too short (mapping length <2/3 of read length) | 8046584.8 (6.90%) | 16700021.9 (14.13%) |
|  | Number of reads unmapped: other | 155314.15 (0.12%) | 146743.7 (0.12%) |
|  | Number of chimeric reads | 0 (0.00%) | 0 (0.00%) |
| Read<br>splices | Number of splices: Total | 95,218,875.7 | 85,817,238.65 |
|  | Number of splices: GT/AG | 93,963,855.4 | 84,565,642.5 |
|  | Number of splices: GC/AG | 1,179,273.45 | 1,184,021.35 |
|  | Number of splices: AT/AC | 75,746.85 | 67,574.8 |
|  | Number of splices: Non-canonical | 0 | 0 |
| Sequence<br>variation | Mismatch rate per base, % | 0.00% | 0.00% |
|  | Deletion rate per base, % | 0.01% | 0.02% |
|  | Deletion average length | 1.853 | 3.123 |
|  | Insertion rate per base | 0.00% | 0.02% |
|  | Insertion average length | 1.3475 | 2.667 |

### Comparison of high-level statistics on sRTD and cRTD

We compared high level statistics of various features, such as genome coverage, genes, transcripts, exons and introns between sRTD and cRTD (Table 2). At the genome level, cRTD covers approximately 91.16 million bases on the genome while sRTD covers 103.79 million bases, a 13.85% increase. This is consistent with the higher number of unmapped reads identified during the STAR mapping step (Table 1). At gene level, sRTD and cRTD include a similar number of genes but sRTD has 8.69% more multi-isoform genes. Protein coding genes with significant hits in the UniProt Plant database (Bateman *et al*., 2021) are also 2.26% higher in sRTD (21,365 in sRTD VS 20,893 in cRTD). Differences at the transcript level are more profound. In spite of having 0.36% fewer genes, the number of transcripts in sRTD increased to 144,872 compared to 128,438 in cRTD, a 12.8% (16,434) increase. The number of protein coding transcripts and those with significant hits in the UniProt Plant database in the sRTD are 15.09% and 13.03% higher, respectively, than in the cRTD and the average protein coding length is 16.43 (4.28%) longer. Transcript diversity in the sRTD also shows 13.19% increase, rising to 2.44 from 2.15 transcripts per gene in cRTD (Table 2 and Supplementary File Figure S1). The sRTD tends to have more and longer exons with the average transcript length in sRTD 12.11% (268.56 bp) longer than in cRTD. Combining the effects of longer transcripts and increased transcript diversity, there is a 26.45% increase of the total transcript length (exonic) in sRTD over cRTD. With significantly less genome coverage, shorter transcripts and fewer protein sequences, it is likely that there are more transcript fragments and incomplete gene models in cRTD than sRTD. Associated with the increased transcript diversity in sRTD we observe an increased number of alternative splicing (AS) events in all event categories. The increase ranges from 21.78% for alternative acceptor sites (A3) to 30.98% for mutually-exclusive exons (MX) (Supplementary File Table S1).

**Table 2:** Basic statistics of the assembled sRTD and cRTD.

| Category | sRTD | cRTD | (sRTD-cRTD)/cRTD |
| --- | --- | --- | --- |
| Genome covered bases | 103,793,515 | 91,164,519 | 13.85% |
| Gene number | 59,447 | 59,664 | -0.36% |
| Multi-isoform gene number | 16,163 | 14,871 | 8.69% |
| Mono-exon gene number | 32,833 | 33,001 | -0.51% |
| Multi-exon gene number | 26,614 | 26,663 | -0.18% |
| <sup>a</sup> Protein-coding | 33,254 (55.94%) | 31,961 (53.57%) | 4.05% |
| <sup>b</sup> Best-hit in UniProt Plant (e < 0.01) | 21,365 (35.94%) | 20,893 (35.02%) | 2.26% |
| Transcript number | 144,872 | 128,438 | 12.80% |
| Mono-exon transcript number | 33,429 | 33,578 | -0.44% |
| Multi-exon transcript number | 111,443 | 94,860 | 17.48% |
| Protein-coding | 73,998 (51.08%) | 64,294 (50.06%) | 15.09% |
| Protein average length | 399.93 | 383.5 | 4.28% |
| Best-hit in UniProt Plant (e < 0.01) | 47,614 (32.87%) | 42,126 (32.80%) | 13.03% |
| Transcript number per gene | 2.44 | 2.15 | 13.19% |
| Transcript N50 | 3,284 | 3,056 | 7.46% |
| Transcript N90 | 1,457 | 1,286 | 13.30% |
| Transcript average length (exonic) | 2,486.78 | 2,218.22 | 12.11% |
| Transcript total length (exonic) | 360,264,709 | 284,903,601 | 26.45% |
| Exon number | 881,447 | 722,816 | 21.95% |
| Exon number per transcript | 6.08 | 5.63 | 8.10% |
| Exon average length | 408.72 | 394.16 | 3.69% |
| Intron number | 736,575 | 594,378 | 23.92% |
| Intron number per transcript | 5.08 | 4.63 | 9.85% |
| Intron average length | 643.8697 | 655.8377 | -1.82% |

<sup>a</sup> Protein-coding genes are defined as those genes with at least one protein-coding transcript.
<sup>b</sup> The number of genes which have at least one protein-coding transcript with the best-hit in the UniProt Plant database.

### Transcript sequence comparisons

We used Blastn to compare the transcript sequences between sRTD and cRTD. To estimate the technical variations caused by Blastn itself, we also compared transcript sequences in sRTD to itself. We defined precision, recall and their weighted mean F1 score to evaluate the proportions of matched bases to the total sequence lengths (Figure 2A). At nucleotide base level, on average 95.2% (precision) of the transcript nucleotides in cRTD are the same as sRTD. Less than 5% of transcript sequences are unique to cRTD. However, 11% of the nucleotide bases from sRTD are missing from the cRTD, indicating a loss of >10% of transcript sequence information when mapping RNA-seq reads to the common reference, consistent with the lower genome coverage identified above.

**Figure 2:**
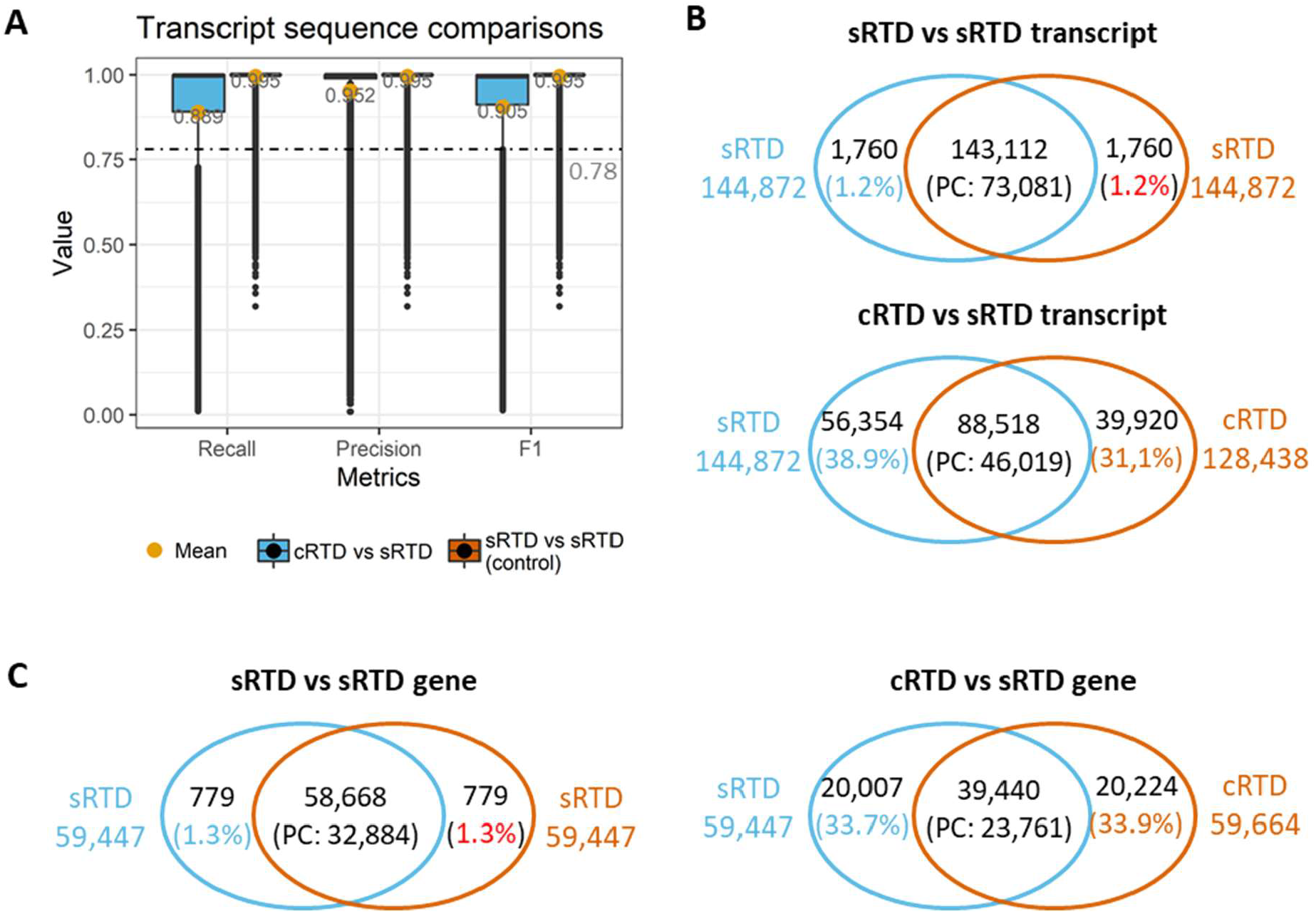
Summary of transcript sequence comparisons of sRTD and cRTD. (A) Metrics distribution of transcript sequence comparisons. The transcript sequences of cRTD and sRTD (query) were both compared to sRTD (target) with Blastn. Recall was defined as number of sequence matched bases over the target sequence length while precision was the matched bases divided by query sequence length. F1 score was the harmonic mean of recall and precision. (B) Venn diagram of transcript-to-transcript match. The overlaps of sRTD vs sRTD and cRTD vs sRTD were those transcripts with sequence similarity F1 > 0.78 (also see Supplementary File Figure S2). PC: protein coding. (C) Gene level comparisons. The overlap represents the genes of matched transcript in (B).

At the individual transcript level, we use an F1 value of 0.78 (the lower limit of outliers of cRTD and sRTD sequence similarity, Figure 2A and Supplementary File Figure S2A) as a cut-off to determine transcripts with a high proportion of sequence overlap. With this threshold, Blastn identified 98.8% of transcripts between sRTD and itself, indicating a low level of technical false positives and negatives (both at 1.2%) caused by Blastn and the chosen parameters. In comparison between cRTD and sRTD, we find that 39,920 (31.1%) transcripts in cRTD do not have a matched transcript of sufficiently high sequence similarity in sRTD. These are likely to be mis-assembled transcripts. 56,354 (38.9%) of the transcripts in the sRTD do not have a corresponding transcript in the cRTD, representing transcripts that failed to assemble when mapping to the common reference (Figure 2B). The transcript sequence discrepancy between cRTD and sRTD directly affects the translated proteins. Only 46,019 out of 73,081 protein coding transcripts in sRTD have a match in cRTD, indicating that 37% of the protein coding transcripts are not assembled or with sufficient similarity to those in sRTD (Figure 2B and Supplementary File Figure S2B). When summarised to the gene level, 20,007 (33.7%) genes in sRTD do not find any genes of sufficiently high transcript sequence similarity in cRTD, while 20,224 (33.9%) genes in cRTD have no corresponding gene models in sRTD. Only 72.3% of the protein coding genes in sRTD have been retrieved in cRTD with at least one matched protein sequence.

### Transcript structure comparisons

To analyse the transcript structure on the same genome coordinate system, we mapped the whole transcript sequences of sRTD and cRTD to the Barke genome with Minimap2, denoting as sRTD.minimap2 and cRTD.minimap2 respectively. Similar to Blastn analysis, the former was used as technical control to estimate the errors introduced by Minimap2. We used Gffcompare to evaluate the structure match of sRTD.minimap2 and cRTD.minimap2 to sRTD at various levels, including nucleotide bases, exon, intron, intron chain, transcripts and gene loci (clusters of exon-overlap). In the Gffcompare analysis, the overlapped gene models shared the same loci. Duplicate transcripts or transcripts whose segments were aligned to multiple chromosomes and/or strands by Minimap2 were discarded, which filtered out 247 transcripts in sRTD.minimap2 and 1,906 transcripts in cRTD.minimap2 (Table 3). The evaluation statistics of true positives was defined as those features that had identical boundaries between the RTDs for comparison (Supplementary File Figure S3). Minimap2 can accurately map the Barke based assembly back to Barke genome with minor technical errors of both false positives and false negatives at < 2% at all levels (Table 3 and 4). However, by using the common reference (Morex) for transcript assembly, we lose ca. 10% introns and exons defined by their genomic co-ordinates and 20% of gene loci. We also have up to 13% of novel predictions that do not exist in the sRTD (Table 3). These observations indicate that the common reference genome poses a significant challenge in studying the genotype specific genes and transcriptional and post transcriptional regulation, including alternative splicing, alternative transcriptional starts as well as polyadenylation site selection.

**Table 3:**
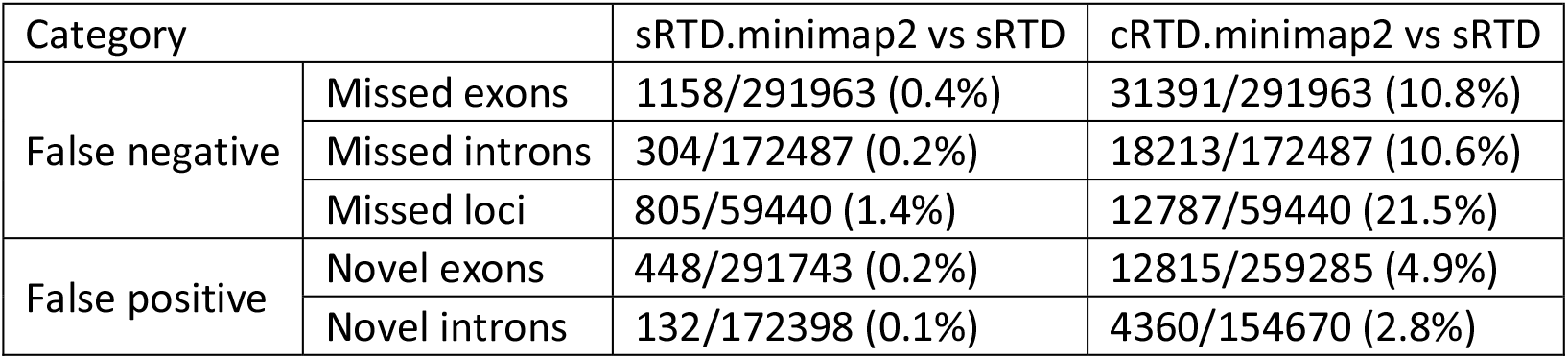
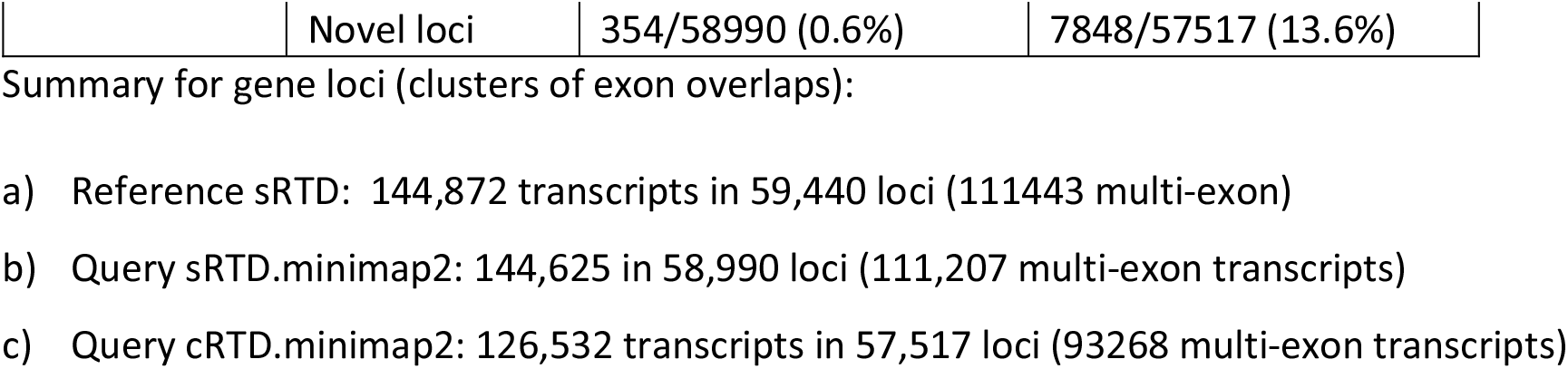
Missed and novel exon, intron and loci regions of Barke genome.

**Table 4:** Recall and precision of structure comparisons at transcript and transcript unit level.

| Category | sRTD.minimap2 vs sRTD |  | cRTD.minimap2 vs sRTD |  |
| --- | --- | --- | --- | --- |
|  | Recall | Precision | Recall | Precision |
| Base level | 99.5% | 99.8% | 76.9% | 90.2% |
| Exon level | 99.3% | 99.4% | 74.1% | 81.4% |
| Intron level | 99.6% | 99.6% | 84.6% | 94.4% |
| Intron chain level | 98.8% | 99.0% | 57.6% | 68.6% |
| Transcript level | 98.6% | 98.6% | 56.8% | 64.6% |
| Locus level | 98.1% | 98.1% | 62.5% | 64.6% |

At the transcript level, Gffcompare classified multi-exon transcripts with identical intron chain or mono-exon transcripts with significant overlap (more than 80% to the longer transcript; default by Gffcompare) as equivalent transcripts. The matched genes must have at least one matched transcript (Supplementary File Figure S2). In Table 4, we can see that cRTD.minimap2 loses about 40% of the total transcripts, multi-isoform transcripts (intron chains) and gene loci present in sRTD, and has about 35% novel assemblies where the structure cannot be found in sRTD, agreeing with the transcript sequence comparisons in Figure 2B. The detailed sub-categories of the transcript structure differences are shown in Supplementary File Figure S3. The largest categories of unmatched cRTD transcripts are fragments with: 1) only one side of a splice junction correctly matched (8.70%); 2) correct splice junctions (7.63%) and 3) retained introns (7.18%). The metrics improve when scaled down to smaller units (nucleotide bases, exons and introns). The introns are the best retrieved, with 85% recall and 94% precision. The matched bases have slightly lower recall (77%) and precision (90%). Due to difficulties in matching the boundaries of first and last exons, the statistics for the exons are the lowest, at 74% and 81%, respectively. The drastic decrease of recall and precision when combining the introns and exons into transcripts is mainly due to the complexity of alternative splicing and variations at the 5’ and 3’ sites.

### Alternative splicing event comparisons

By using the Minimap2 results, we can directly compare the AS events between cRTD.minimap2 and sRTD.minimap2 on their genomic coordinates. We used SUPPA2 to generate local AS events of retained intron (RI), alternative 5’ splice-site (A5), alternative 3’ splice-site (A3), skipping exon (SE), alternative first exon (AF), alternative last exon (AL) and mutually exclusive exons (MX)at the splice junctions of multi-exon transcripts (Trincado *et al*., 2018). The AS events of cRTD.minimap2 and sRTD.minimap2 are treated as queries and compared to the target events in sRTD (Table 5). The comparisons of sRTD.minimap2 and sRTD revealed minor technical errors for all the events, with Precision and Recall both close to 1. The precision and recall of cRTD.minimap2 against sRTD varies at different events. The A3, A5 and SE events are the best matched, but still about 10% of splice junctions failed to be identified, and we observed a 30% false discovery rate. The majority of AF and AL events in cRTD.minimpa2 are incorrect predictions, which is consistent with our earlier analysis that cRTD tends contains short fragmented transcripts, which are likely to be mis-annotated at the transcript ends.

**Table 5:** Comparisons of alternative splicing events.

| Comparisons | AS events | overlaps between query and targets |  |  | Precision | Recall |
| --- | --- | --- | --- | --- | --- | --- |
|  |  | Query | Query & Target | Target |  |  |
| sRTD.minimap2 (Query)<br>vs sRTD (Target) | RI | 216 | 32351 | 335 | 0.990 | 0.993 |
|  | A3 | 94 | 22899 | 72 | 0.997 | 0.996 |
|  | A5 | 83 | 16552 | 51 | 0.997 | 0.995 |
|  | SE | 30 | 12973 | 74 | 0.994 | 0.998 |
|  | AF | 7 | 1666 | 11 | 0.993 | 0.996 |
|  | AL | 15 | 1488 | 17 | 0.989 | 0.990 |
|  | MX | 9 | 743 | 18 | 0.976 | 0.988 |
| cRTD.minimap2 (Query)<br>vs sRTD (Target) | RI | 9626 | 15906 | 16780 | 0.487 | 0.623 |
|  | A3 | 2345 | 16282 | 6689 | 0.709 | 0.874 |
|  | A5 | 1748 | 11462 | 5141 | 0.690 | 0.868 |
|  | SE | 1195 | 8765 | 4282 | 0.672 | 0.880 |
|  | AF | 870 | 384 | 1293 | 0.229 | 0.306 |
|  | AL | 639 | 515 | 990 | 0.342 | 0.446 |
|  | MX | 111 | 441 | 320 | 0.580 | 0.799 |

The logical relation between sets illustrates the unique and overlap sets of Query and Target. Precision: (Query & Target)/Query; Recall: (Query & Target)/Target

We further studied the percentage spliced-in (PSI) of sRTD.minimap2, cRTD.minimap2 and sRTD, a measure of the relative abundance of AS events. We used Salmon and the Barke RNA-seq reads of 20 samples to generate TPMs for event PSI calculation in SUPPA2. We find that the Pearson and Spearman correlations between PSI values in cRTD.minimap2 and identical events in sRTD were significantly lower than that in the control (comparison of sRTD.minimap2 and sRTD) for all the AS events, while the mean relative errors (absolute value of PSI difference divided by PSI of sRTD) of 20 samples were significantly higher (Figure 3). Thus, transcript assembly from the cRTD is particularly problematic when used for alternative splicing analysis.

**Figure 3:**
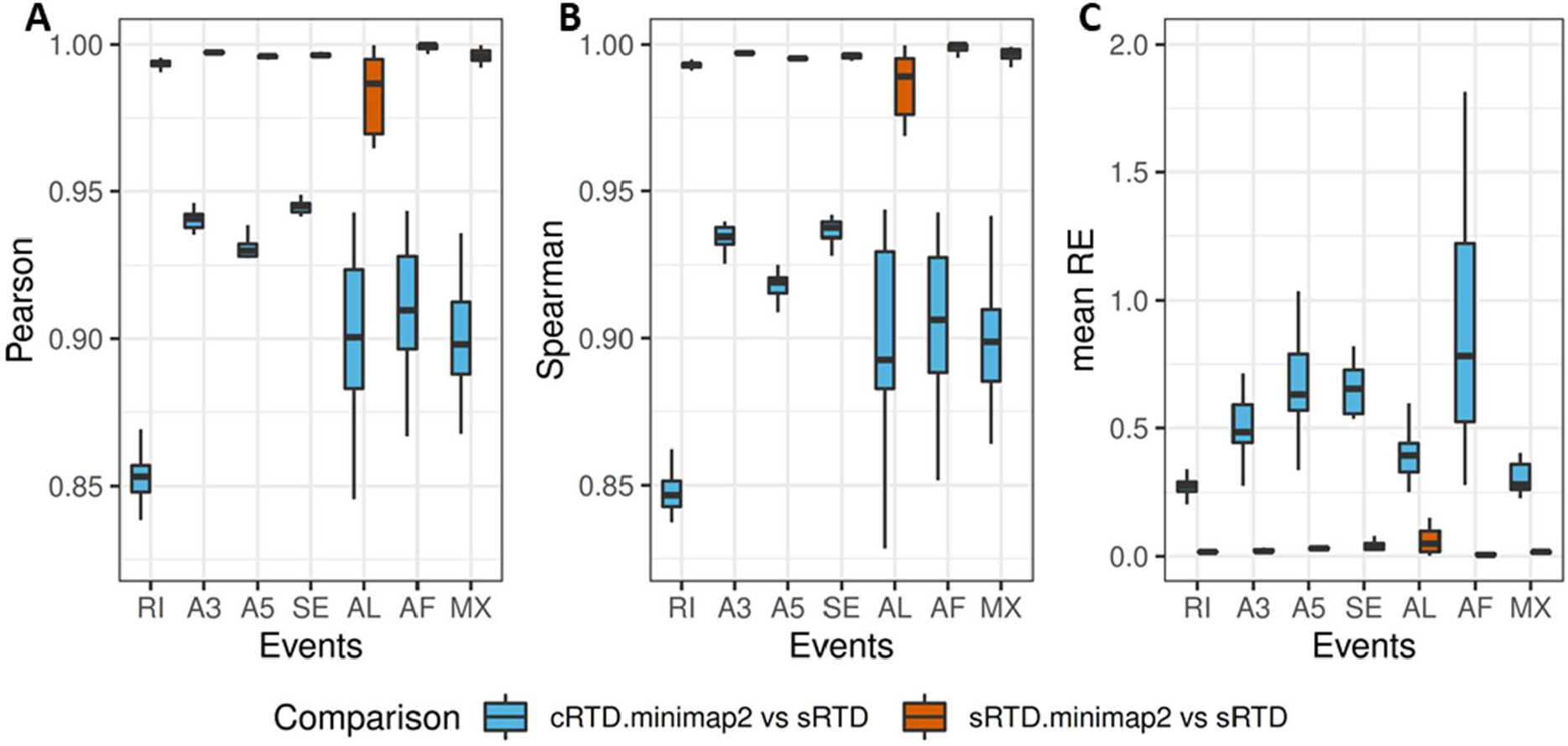
Comparisons of AS analysis accuracy. The Pearson correlation (A), Spearman correlations (B) and mean relative errors (C) were calculated for different AS events in 20 samples. The outliers of the distributions were removed. RI: retained intron, A5: alternative 5’ splice-site, A3: alternative 3’ splice-site, SE: skipping exon, AF: alternative first exon, AL: alternative last exon and MX: mutually exclusive exons.

### Transcript quantification comparisons

To investigate how the expression estimation for individual genes are affected and what the characteristics are of the genes that are most affected, we generated simulated RNA-seq reads to assess the quantification errors introduced by using different RTDs. We used the read count matrix generated from Barke caryopsis and root tissues (each with 3 biological reps) to simulate the RNA-seq data using the Polyester R package based on sRTD (Frazee *et al*., 2015). The corresponding transcripts between cRTD and sRTD were obtained using the Blastn sequence match with F1 score > 0.78 and Gffcompare match with identical transcript structures (Supplementary File Table S2). Thus, the transcript quantifications of cRTD and sRTD can be directly compared to the read count matrix (ground truth) used for the simulation. Although we used stringent criteria to define the equivalent transcripts, we find that the Pearson and Spearman correlations of comparisons between sRTD and ground truth are constantly higher than that of cRTD and ground truth (Table 6). The relative errors are significantly higher between cRTD and ground truth at both gene and transcript level (Figure 4). sRTD outperforms cRTD in all categories at both transcript and gene level; The relative errors are 2 to 3 fold higher when using cRTD for quantification even at the gene level, indicating sRTD provides more accurate quantification at both transcript and gene level. The results also show that quantification is more accurate at gene level than at the transcript level for the RNA-seq data.

**Table 6:** Average correlation of transcript quantification comparisons.

| Quantification | Correlation | sRTD vs Truth | cRTD vs Truth |
| --- | --- | --- | --- |
| Gene level | Pearson | 0.9996 | 0.9966 |
|  | Spearman | 0.9934 | 0.9865 |
| Transcript level | Pearson | 0.9993 | 0.9824 |
|  | Spearman | 0.9242 | 0.8804 |

**Figure 4:**
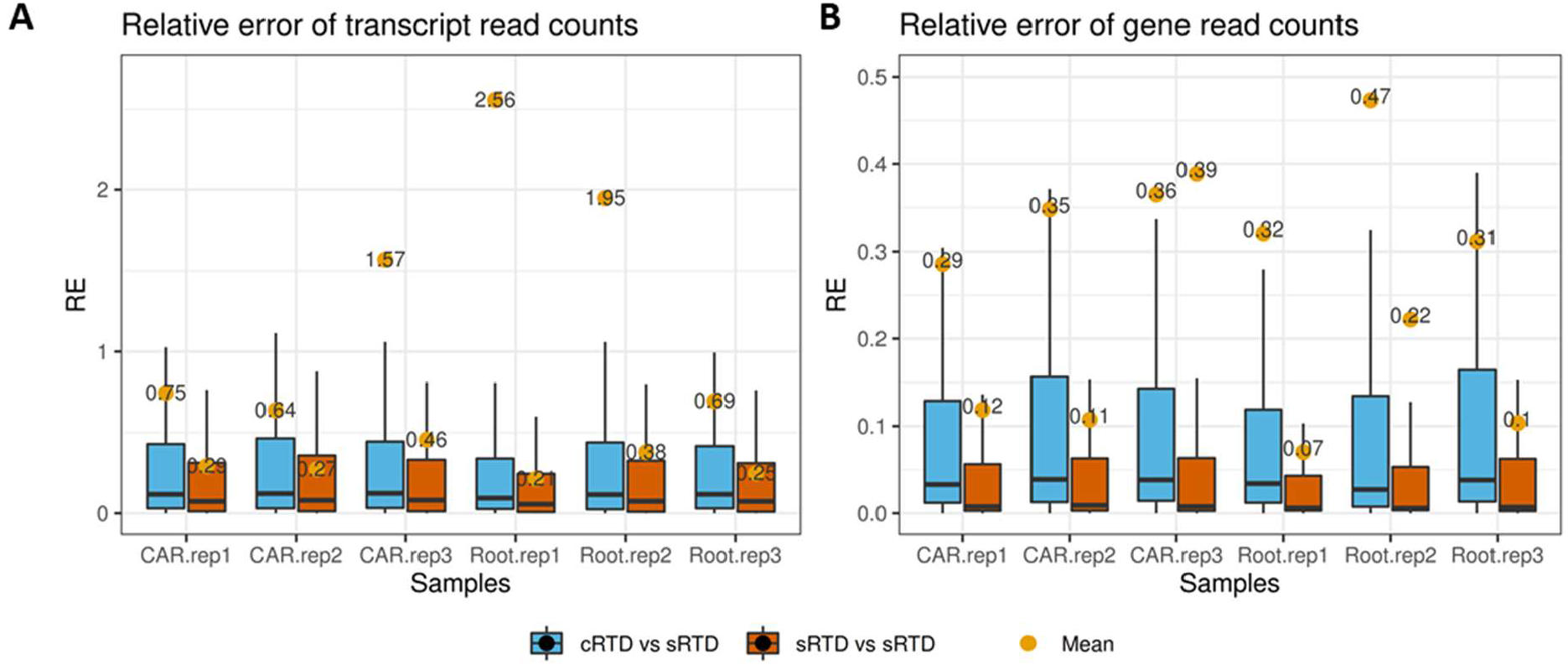
Distribution of read count relative errors. The read counts from sRTD and cRTD quantifications were both compared to ground truth for relative error calculation at (A) transcript level and (B) gene level. The outliers of were removed and the dots highlighted the mean values of the distributions.

The cRTD transcripts with lower similarities to sRTD (i.e. F1 score < 0.78) were also investigated in terms of quantification accuracy. The transcripts in cRTD were classified into eight groups according to their F1 scores in intervals of 0.1 between 0-0.78 (Figure 5). The relative error of the quantification increases from 0.5 (50%) to 1 (100%) as the sequence similarities between corresponding transcript in cRTD and sRTD decreases. The average correlation between the transcript abundance estimation using cRTD and the ground truth also improves (Pearson correlation from 0.367 to 0.81, Spearman correlation from 0.028 to 0.65) with an increasing similarity score. Thus, transcripts in cRTD with sequences different from those in sRTD (i.e. 38.9% of transcripts with F1 score <0.78) would produce significant quantification errors and should therefore be a cause of concern when used for quantification and differential expression analysis.

**Figure 5:**
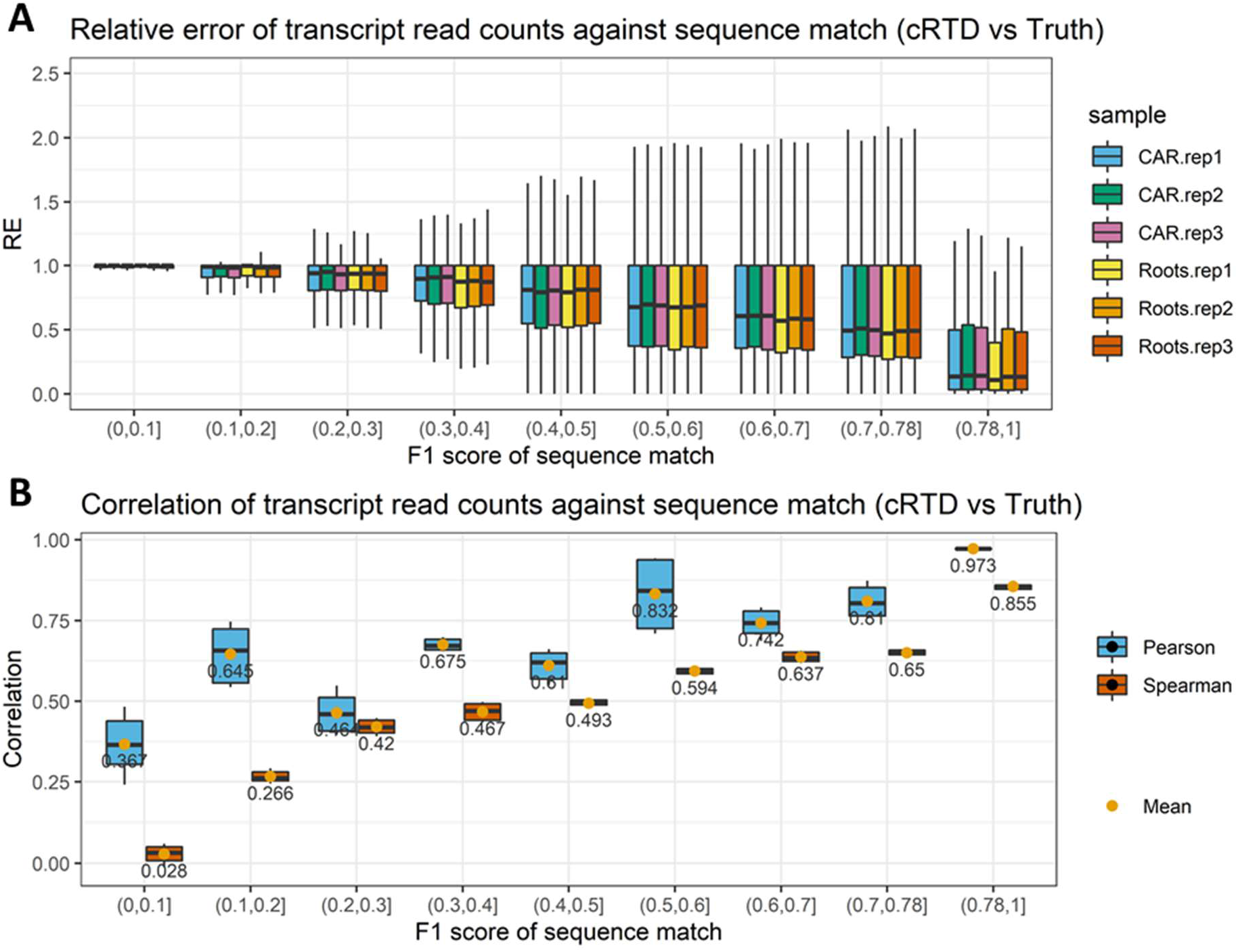
Comparison of transcript quantification of cRTD and ground truth at different sequence similarities. F1 scores of sequence match using Blastn are divided into different intervals. The (A) relative errors and (B) Pearson and Spearman correlations of read counts are calculated for the matched transcripts within corresponding F1 score intervals. The outliers of the boxplots are removed.

### Differential expression and alternative splicing

We further investigated the downstream impact on differential expression analysis with the simulated RNA-seq data. We used the 3D RNA-seq pipeline to investigate 1) differentially expressed (DE) genes and transcripts; 2) differentially alternatively spliced (DAS) genes; and 3) differential transcript usage (DTU) transcripts between the caryopsis and roots tissues. The PCA plots (Supplementary File Figure S4) reveal that the quantifications obtained using both sRTD and cRTD successfully capture the variation in the data between caryopsis and root tissues as well as the biological replicates of ground truth. The amount of the variation explained by PC1 and PC2 in all three datasets are not significantly different. Comparisons between DE genes, DAS genes, DE transcripts and DTU transcripts illustrate that DE analysis at the gene level is most stable and least affected by the choice of RTD. sRTD shows precision and recall both >99% for DE genes, while the cRTD only achieves 67.7% precision and 67.0% recall, missing about 4000 DE genes while inferring 4000 false positive genes. The transcript level analysis is more variable, but the analysis for DAS genes and DTU transcripts both shows the superior performance of sRTD over cRTD in both precision and recall, consistent with our observations that use of the cRTD presents significant challenges for transcriptional level analysis.

We then compared the log_2_ fold change (L2FC) differences of matched genes and transcripts using the quantification results obtained by using two RTDs and the ground truth. The L2FCs at gene level are much closer to ground truth, with greater than one magnitude lower relative errors compared to the transcript level (Figure 6A). When comparing sRTD and cRTD to ground truth, we identified that 1,849 (5.84%) expressed transcripts in sRTD and 2,276 (7.19%) in cRTD have inverted up-and down-regulation (Figure 6B). At the gene level, the number of inversions significant decrease to 103 (0.58%) and 268 (1.51%) for sRTD and cRTD respectively (Figure 6C). Applying a cut-off |L2FC| > 1, the numbers of regulatory switches are further reduced. For example, only two genes in cRTD and one gene in sRTD switch to up-regulation and one gene in cRTD to down-regulation.

**Figure 6:**
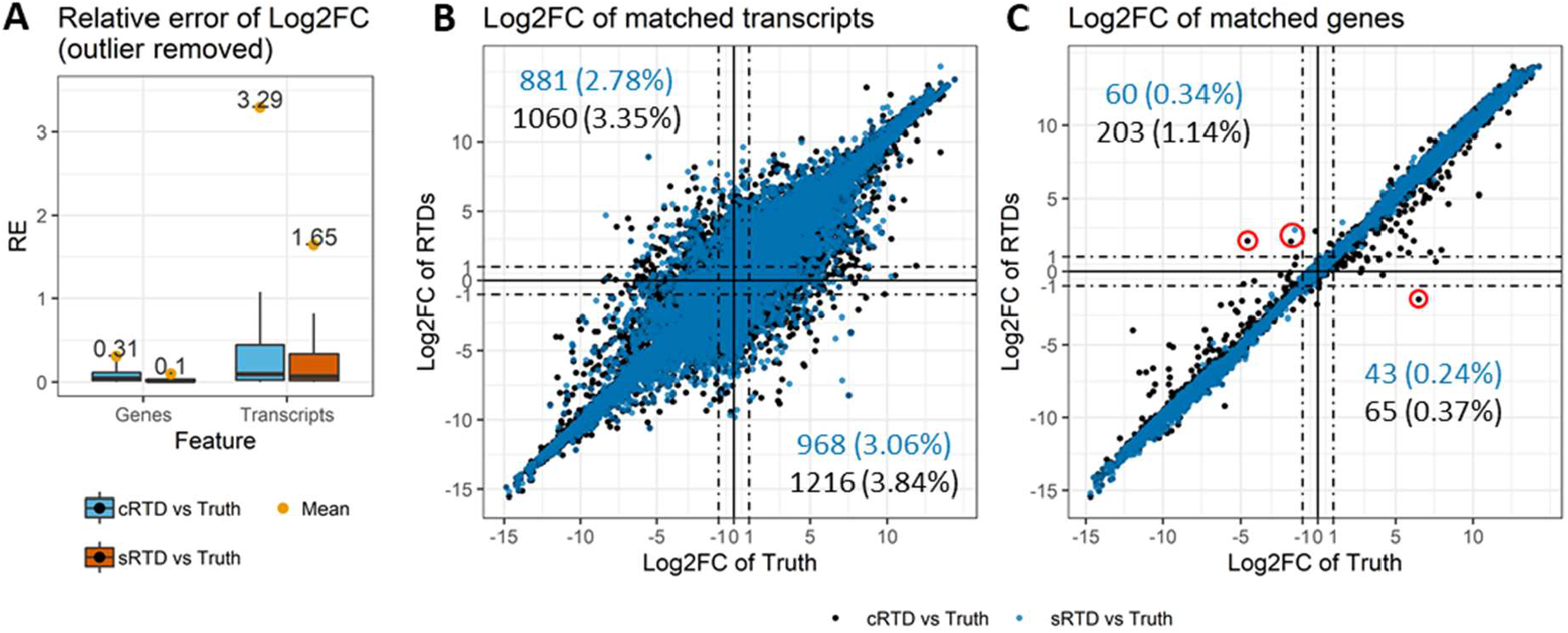
Comparisons of L2FC. The genes and transcripts between cRTD and sRTD were determined by the Blastn sequence and Gffcompare structure analyses. The low expressed transcripts and genes in Ground truth, sRTD or cRTD were filtered, yielding 31,640 transcripts and 17,768 genes. (A) L2FC relative error distributions of matched genes and transcripts. (B) L2FC of matched transcripts. (C) L2FC of matched genes. In (B) and (C), the numbers and percentages on the top-left and bottom-right quadrants demonstrate the transcripts and genes that switch up- and down-regulations. The dashed lines highlight the L2FC cut-offs of 1 and -1 that we often use to determine the change significance. The red circles in (C) highlight the genes that switch up- and down-regulation with |L2FC| ≥ 1.

## Discussion

In this study, we quantitatively evaluated the impacts of using a common reference genome versus a genotype specific genome for transcriptome analysis. We used data from two cultivated barley genotypes, the American cultivar Morex and the European cultivar Barke, both with available high quality genome assemblies. We mapped Barke RNA-seq reads to both Morex and Barke genomes to assemble a common reference-based cRTD (Morex) and a genotype-specific sRTD (Barke) respectively. We used same parameters for read mapping, transcript assembly and various comparisons throughout. By setting the reference genome as the only variable and comparing the assembled sRTD to itself to estimate inherent technical variation, we were able to investigate the impacts of using a common reference on the outcomes of transcriptomic data analyses from different but related genotypes.

Although the assembled gene numbers were comparable in sRTD and cRTD, the use of a common reference genome resulted in a significant reduction in transcript diversity. Read alignment to the Morex genome produced 11.6% fewer splice junctions which led to a >20% loss of introns and exons in the assembled cRTD. The reduction of splice junctions also resulted in >20% fewer alternative splicing events as well as >10% less transcript isoforms. We found that the transcripts in cRTD are 268.56 nucleotides shorter on average. This indicates that many transcripts assembled using a common reference genome are likely to be fragmentary and incomplete. We found that the reduced diversity and incomplete transcript assemblies in the cRTD reduce the number of protein-coding transcripts by 15% and protein coding genes by 4%. The average protein sequence length is also 16.43 amino acids shorter implying that the fragmentation and incompleteness of cRTD transcripts affects their coding capacity.

At the individual transcript sequence level, we used Blastn to study the sequence similarity between cRTD and sRTD. Blastn locally aligns stretches of sequences with a high level of base matches without considering diverging regions (Altschul *et al*., 1990). Thus, two transcript sequences may have significant Blastn e-value and high bit-score, but have only a small proportion of overlap. Thus, we defined an “F1 score” to identify transcript pairs with significant sequence overlap proportional to the full length of sequences. We also used Minimap2 to align the cRTD transcript to the Barke genome and Gffcompare to identify the transcripts with identical structures to those in sRTD. By combining the sequence and structure comparisons, we identified 78,540 (61.2%) transcripts in cRTD that find an equivalent transcript in sRTD with significant sequence as well as structure similarity (Supplementary File Table S2). We also observed disagreements between sequence and structure matches. We identified 1,776 transcripts that had identical structure in sRTD and cRTD, but with significantly different sizes of first and/or last exons, thereby leading to low sequence similarity of F1 ≤ 0.78 (Supplementary File Table S2 and Figure S5A). On the other hand, 9,978 transcripts with high sequence similarity (i.e. F1 > 0.78) had distinct structures, which were due to inconsistent intron sizes, such as shift of intron boundaries, missing and novel introns caused by technical errors in Minimap2, or sequence variation arising from mapping Morex-based assemblies to the Barke genome (Supplementary File Table S2 and Figure S5B-D). Thus the statistics for sequence and structure comparisons shows different level of similarities, especially for the nucleotide base level (Figure 2 and Table 4).

Allowed mismatches is a key parameter affecting the number of reads mapped to a reference genome and makes a significant impact on transcript assembly. We explored the influence of this parameter by allowing 0, 2, 4 and 6 mismatches in the second pass of read mapping during sRTD and cRTD assembly (Supplementary File Table S3-5). We observed a general trade-off in that by allowing more mismatches, more reads are mapped but the chances of mis-mapping also increase. When mapping Barke RNA-seq reads to the Morex genome allowing 2 mismatches, 107,619,513 (89.78%) reads map uniquely, compared to 107,770,451.15 (89.89%) when mapping the same data to the Barke genome allowing 0 mismatches. We therefore used Gffcompare to assess differences in the transcript structures in a cRTD constructed allowing 2 mismatches, denoted as cRTD_m2 (Supplementary File Table S6). When allowing 2 mismatches, the number of missed gene loci in cRTD_m2 reduced by 1,979 from 21.5% to 18.2%. However, the number of false positive loci increased by 3,600 from 13.6% to 18.6%. Thus, allowing 2 mismatches enables more genes to be discovered but this is accompanied by more false discoveries. For individual transcripts, we observe decreased precision at almost all levels when allowing mismatches, whereas recall increases at almost all levels. Thus, to focus on the impact of genome reference and to minimize the confounding of mismatch settings, we used a cRTD constructed with the parameter setting of 0 mismatch for all the quantitative analysis. The errors and false discovery rates we use are therefore conservative estimates.

Using the sRTD improved the quantifications of transcript abundances and downstream differential expression analysis. We used simulated RNA-seq read data to compare transcript quantifications revealed using cRTD and sRTD. We found that the quantifications of transcripts in sRTD have higher correlation and lower errors to the ground truth. Not surprisingly, the quantification error increases and correlation decreases as the transcript sequence discrepancies increase between cRTD and sRTD (Figure 5). However, gene level quantification of transcript abundance appears to be much more robust and less affected by the RTD with the relative error almost one magnitude lower compared to transcript level quantifications. Quantification accuracy also directly affects the downstream differential expression analysis. Our comparisons indicate that with stringent filters of significance, only around 70% of the DE genes quantified using cRTD agree with ground truth (Table 7). The error of L2FC at gene level is over one magnitude smaller than at transcript level. The error of L2FC is significantly higher and we have fewer correct identifications of DE transcripts, especially the alternative splicing related DAS genes and DTU transcripts (Table 7). Finally, there are only a few outlier genes that appear to switch between up-and down-regulation when assessed using the different RTDs (change sign of L2FC). The number gets further reduced when applying additional filters, for example |L2FC|≥1 (Figure 6).

**Table 7:** Comparisons of differentially expressed and alternative spliced genes and transcripts.

| Category | Comparison | Overlaps between Query and Target |  |  | Precision | Recall |
| --- | --- | --- | --- | --- | --- | --- |
|  |  | Query | Target & Query | Target |  |  |
| DE genes | sRTD vs Truth | 117 | 12047 | 122 | 0.990 | 0.990 |
|  | cRTD vs Truth | 3893 | 8149 | 4020 | 0.677 | 0.670 |
| DAS genes | sRTD vs Truth | 77 | 298 | 188 | 0.795 | 0.613 |
|  | cRTD vs Truth | 111 | 160 | 326 | 0.590 | 0.329 |
| DE transcripts | sRTD vs Truth | 776 | 10098 | 1391 | 0.929 | 0.879 |
|  | cRTD vs Truth | 3903 | 6777 | 4712 | 0.635 | 0.590 |
| DTU transcripts | sRTD vs Truth | 181 | 499 | 337 | 0.734 | 0.597 |
|  | cRTD vs Truth | 222 | 272 | 564 | 0.551 | 0.325 |

- The unique and overlap sets in the comparisons between sRTD/cRTD (Query) and ground Truth (Target). Precision: (Query & Target)/Query; Recall: (Query & Target)/Target. - The overlap sets represent the matched transcripts between sRTD/cRTD and Ground Truth with criteria F1 > 0.78 in Blastn sequence match and identical structure in Gffcompare match. - The matched gene must have at least one matched transcript.

In summary, due to the difficulty in assembling transcripts accurately using short reads, transcript level analysis remains challenging. When using contemporary pseudoalignment methods for assessing transcript abundance the impact of the source RTD on quantification accuracy can be considerable for transcript level analysis. However, reassuringly for the majority of transcriptomic analyses, working at gene level appears relatively robust and represents a reasonable, if potentially less informative, compromise when a high quality and comprehensive genotype specific transcript reference is not available.

## Data availability

The RNA-seq data used in this study is available at SRA with BioProject accession number PRJNA755523. The core scripts of the data analysis can be viewed from: https://github.com/wyguo/genotype_specific_RTD.

## Acknowledgements

Author Contributions: R.Z. supervised and managed the project. W.G. designed the pipeline and performed the whole analysis. R.W and M.C. generated part of the barley RNA-seq data and contributed to the design of the analysis. W.G. and R.Z. integrated the results and initialised the manuscript. All authors read the manuscript and contribute to the finish of the manuscript.

Great thanks also go to Nils Stein and Ronja Wonneberger from Leibniz Institute of Plant Genetics and Crop Plant Research (IPK), Germany for providing part of the RNA-seq data.

