## Supplementary material for "The value of genotype-specific reference for transcriptome analyses": supp materials

### Supplementary File

#### Supplementary Figures

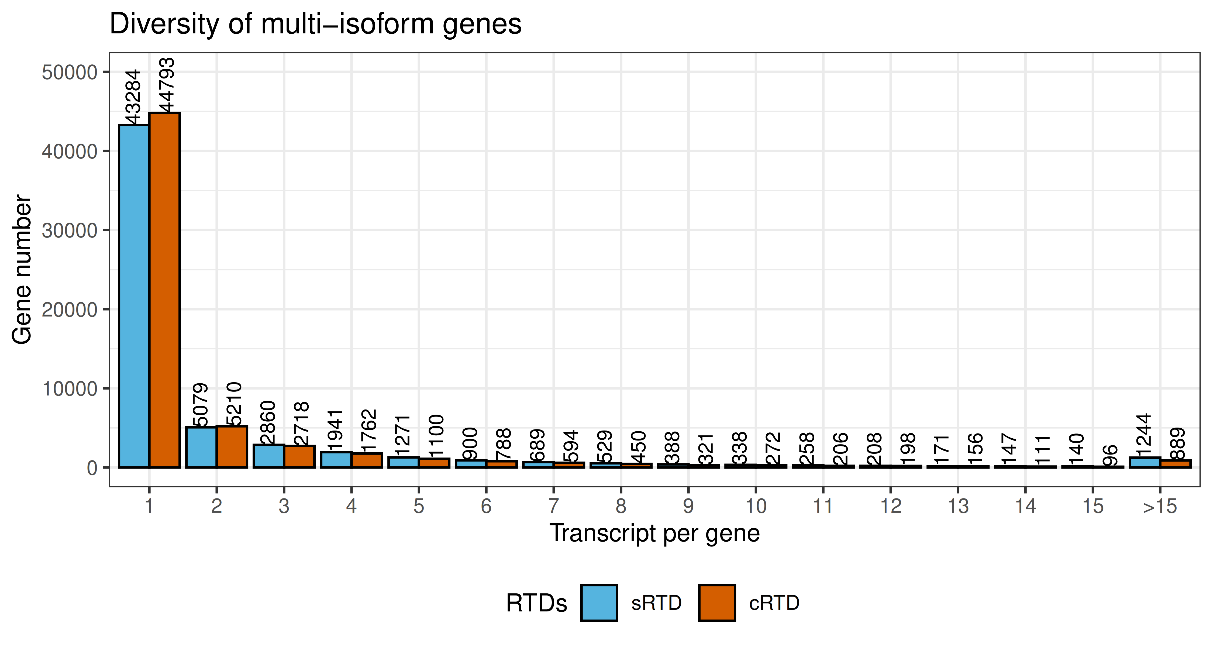

Figure S1: Number of multi-isoform genes in sRTD and cRTD.

X-axis shows the number of transcript isoforms in the genes and y-axis counts the number of genes with that amount of transcript isoforms.

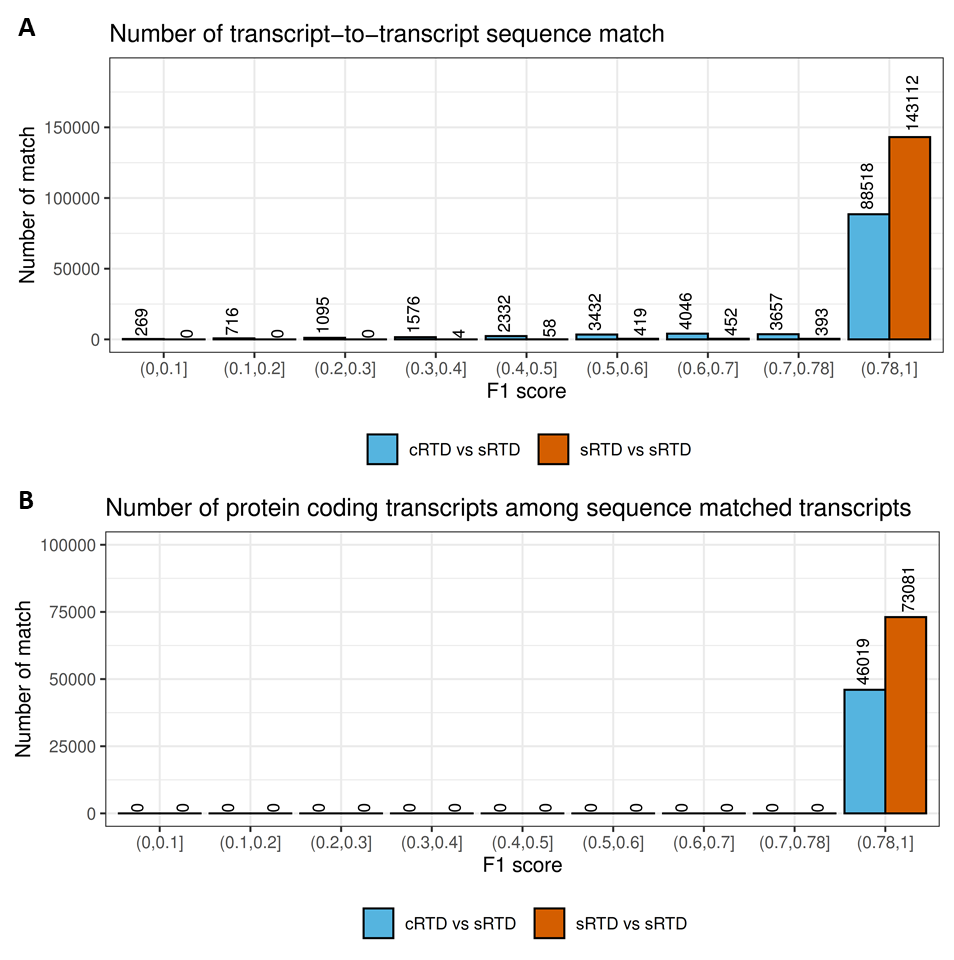

Figure S2: Transcript-to-transcript sequence match results using Blastn.

(A) Number of transcript-to-transcript sequence matches in different intervals of F1 score. The matches between RTDs were classified into different intervals according to their F1 score of sequence similarity. The numbers of matches in each interval are shown on the top of the bars. For example, if the sequence comparison of transcript.1 in cRTD and transcript.2 in sRTD has F1=0.95, we define transcript.1 matches to transcript.2 in F1 interval (0.9,1]. The numbers in the dashed box match to the numbers in the overlaps of the Venn diagram in Figure 1 of main text. (B) The number of matched transcripts with identical protein sequences.

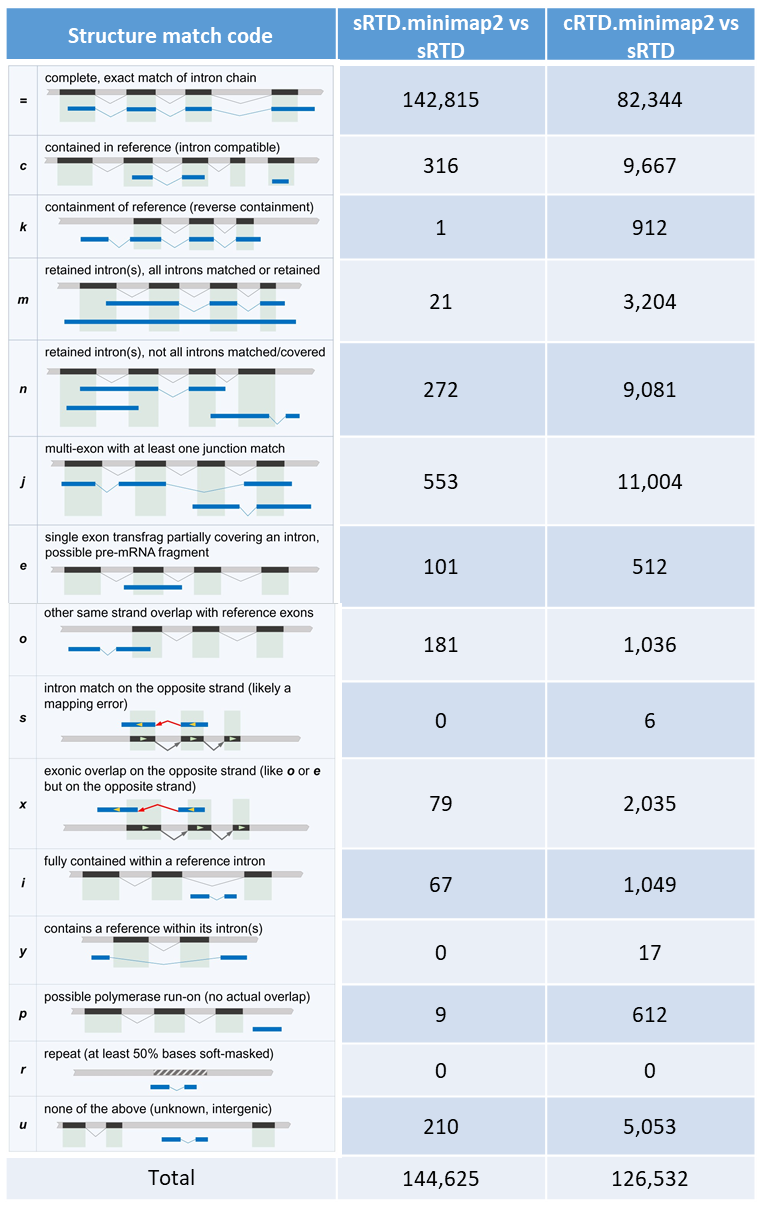

Figure S3: Transcript match codes generated by Gffcompare.

The schema of transcript match classifications were taken from Pertea and Pertea (2020). The numbers in different code classifications were generated by comparing the transcripts of sRTD.minimap2 and cRTD.minimap2 to sRTD, respectively. For example, the match code “=” in the table indicates an exact match of intron chain (combination of introns of a transcript).

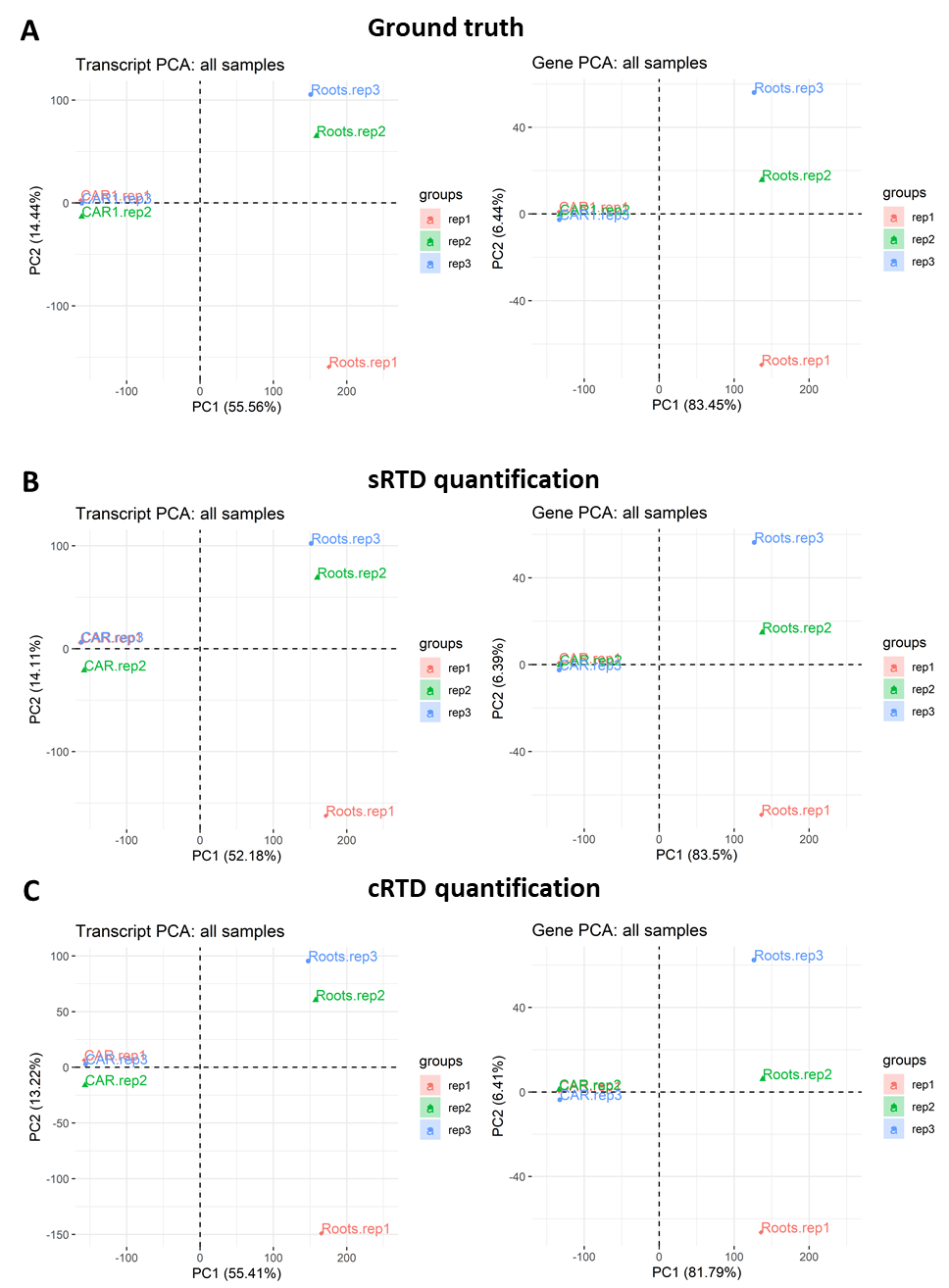

Figure S4: PCA plot of transcript and gene level quantifications.

The PCA plots are based on the quantifications of (A) Ground Truth, (B) sRTD and (C) cRTD.

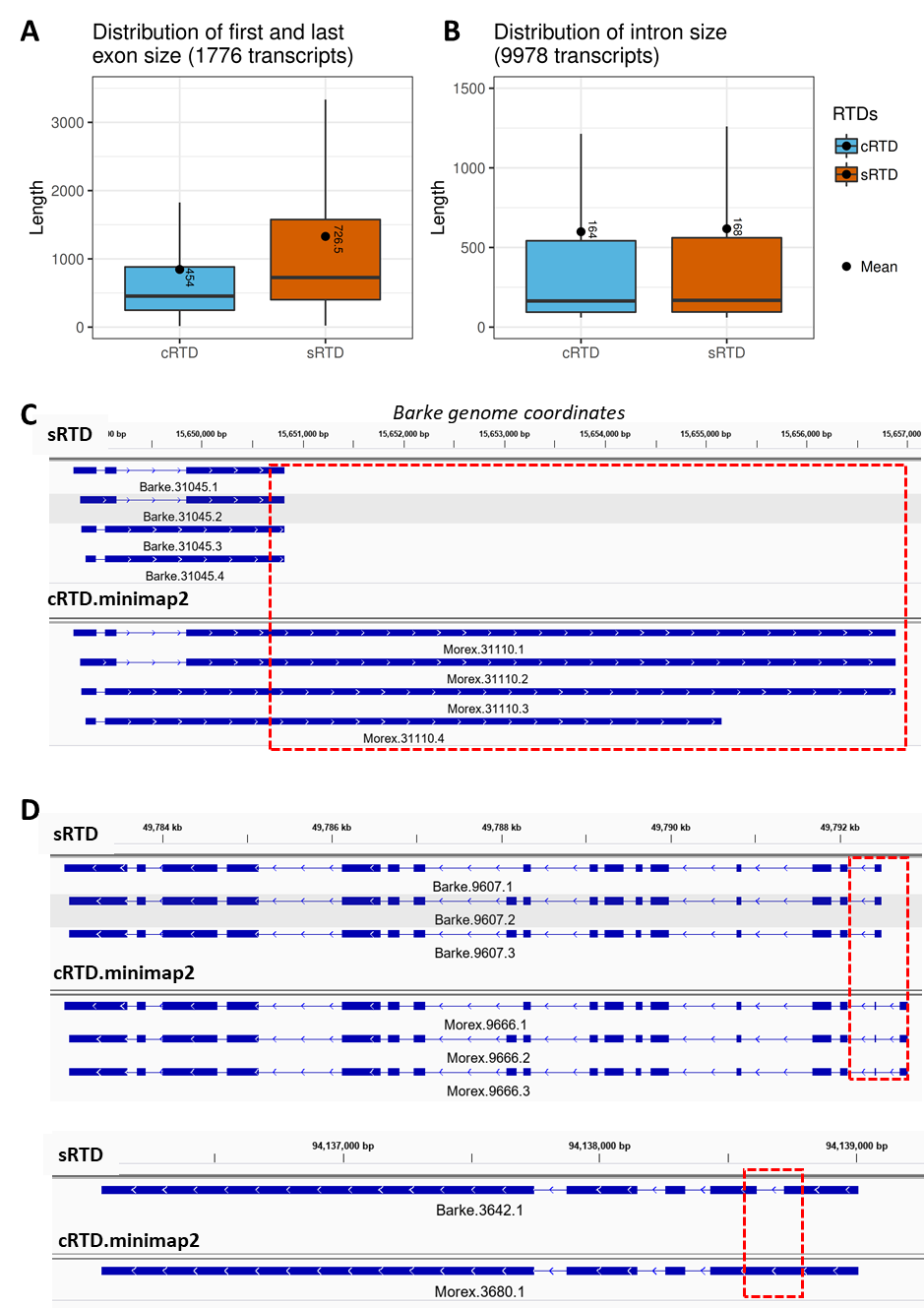

Figure S5: Exon and intron size of inconsistent transcript between sequence and structure match.

(A) Distribution of the first and last exon sizes of 1,776 transcripts with identical structure but low sequence similarity. (B) Distribution of all intron sizes of 9,978 transcripts with high sequence similarity but distinct structures. (C) An example of transcripts in category (A) and (D) Two examples of transcripts in category (B).

#### Supplementary Tables

Table S1: Alternative splicing events of sRTD and cRTD.

| AS events | sRTD | cRTD | (sRTD-cRTD)/cRTD |
| --- | --- | --- | --- |
| RI | 32,686 | 26,409 | 23.77% |
| A3 | 22,971 | 18,863 | 21.78% |
| A5 | 16,603 | 13,358 | 24.29% |
| SE | 13,047 | 10,211 | 27.77% |
| AL | 1,505 | 1,204 | 25.00% |
| AF | 1,677 | 1,268 | 32.26% |
| MX | 761 | 581 | 30.98% |

RI: Retained intron; A3: Alternative 3’ splice-site; A5: Alternative 5’ splice site; SE: Skipping exon; AL: Alternative last exon; AF: Alternative first exon; MX: Mutually exclusive exons.

Table S2: The relation between sequence match and structure match between cRTD and sRTD.

| Comparison | F1 Score | = | c | k | m | n | j | e | o | s | x | i | y | p | miss |
| --- | --- | --- | --- | --- | --- | --- | --- | --- | --- | --- | --- | --- | --- | --- | --- |
| sRTD vs sRTD | (0,0.1] | 0 | 0 | 0 | 0 | 0 | 0 | 0 | 0 | 0 | 0 | 0 | 0 | 0 | 0 |
|  | (0.1,0.2] | 0 | 0 | 0 | 0 | 0 | 0 | 0 | 0 | 0 | 0 | 0 | 0 | 0 | 0 |
|  | (0.2,0.3] | 0 | 0 | 0 | 0 | 0 | 0 | 0 | 0 | 0 | 0 | 0 | 0 | 0 | 0 |
|  | (0.3,0.4] | 3 | 0 | 0 | 0 | 0 | 0 | 0 | 1 | 0 | 0 | 0 | 0 | 0 | 0 |
|  | (0.4,0.5] | 43 | 2 | 0 | 0 | 0 | 0 | 0 | 10 | 0 | 0 | 0 | 0 | 0 | 3 |
|  | (0.5,0.6] | 330 | 0 | 0 | 2 | 0 | 3 | 0 | 22 | 0 | 0 | 0 | 0 | 0 | 62 |
|  | (0.6,0.7] | 365 | 0 | 0 | 3 | 0 | 0 | 0 | 9 | 0 | 0 | 0 | 0 | 0 | 75 |
|  | (0.7,0.78] | 321 | 1 | 0 | 3 | 0 | 4 | 0 | 9 | 0 | 0 | 0 | 0 | 0 | 55 |
|  | **(0.78,1]** | **141142** | 42 | 0 | 7 | 1 | 166 | 74 | 97 | 0 | 0 | 0 | 0 | 0 | 1583 |
| cRTD vs sRTD | (0,0.1] | 1 | 34 | 0 | 0 | 1 | 4 | 4 | 5 | 0 | 5 | 0 | 0 | 0 | 215 |
|  | (0.1,0.2] | 6 | 131 | 1 | 1 | 7 | 28 | 6 | 12 | 0 | 25 | 0 | 0 | 1 | 498 |
|  | (0.2,0.3] | 22 | 230 | 5 | 13 | 17 | 55 | 10 | 24 | 0 | 45 | 0 | 0 | 0 | 674 |
|  | (0.3,0.4] | 57 | 360 | 5 | 36 | 54 | 69 | 10 | 31 | 0 | 61 | 0 | 1 | 2 | 890 |
|  | (0.4,0.5] | 118 | 516 | 14 | 107 | 78 | 100 | 20 | 44 | 0 | 74 | 0 | 0 | 0 | 1261 |
|  | (0.5,0.6] | 332 | 658 | 34 | 182 | 57 | 153 | 24 | 69 | 0 | 82 | 1 | 0 | 0 | 1840 |
|  | (0.6,0.7] | 539 | 860 | 41 | 170 | 73 | 209 | 30 | 64 | 0 | 86 | 0 | 0 | 1 | 1973 |
|  | (0.7,0.78] | 701 | 907 | 53 | 142 | 24 | 160 | 17 | 50 | 1 | 84 | 0 | 0 | 0 | 1518 |
|  | **(0.78,1]** | **78540** | 1930 | 317 | 202 | 20 | 577 | 27 | 101 | 0 | 298 | 1 | 0 | 0 | 6505 |

a) Blastn was used to compare the transcript sequence of sRTD vs sRTD and cRTD vs sRTD. F1 score, which is the weighted average of sequence match recall and precision, was divided into intervals from 0 to 1.
b) Gffcompare was used to compare the transcript structure of sRTD.minimap2 vs sRTD and cRTD.minimap2 vs sRTD. The details of transcript match code in the column names can be found in Figure S3. The last column “miss” indicates the structure match information is missing.
c) The number of matched transcripts with identical intron chains and sequence similarity of F1 > 0.78 were highlighted with bold text. The match columns with 0 transcripts are excluded.

Table S3: STAR second pass mapping statistics at different mismatch settings.

| **Barke-ref mapping (allowed max mismatch)** | **Barke (m=0)** | **Barke (m=2)** | | **Barke (m=4)** | | | **Barke (m=6)** |
| --- | --- | --- | --- | --- | --- | --- | --- |
| Number of input reads | 119,777,808.4 | 119,777,808.4 | | 119,777,808.4 | | | 119,777,808.4 |
| Average input read length | 298.25 | 298.25 | | 298.25 | | | 298.25 |
| Uniquely mapped reads number | 107,770,451.15 (89.89%) | 110,436,414.35 (92.19%) | | 111,149,168.6 (92.82%) | | | 111,532,732.15 (93.15%) |
| Average mapped length | 288.64 | 294.906 | | 295.6715 | | | 296 |
| Number of splices: Total | 95,218,875.7 | 99,597,282.1 | | 100,420,405.7 | | | 100,780,977.3 |
| Number of splices: GT/AG | 93,963,855.4 | 98,287,080.7 | | 99,096,506.45 | | | 99,451,054.9 |
| Number of splices: GC/AG | 1,179,273.45 | 1,231,470.6 | | 1,244,507.75 | | | 1,250,205.55 |
| Number of splices: AT/AC | 75,746.85 | 78,730.8 | | 79,391.45 | | | 79,716.85 |
| Number of splices: Non-canonical | 0 | 0 | | 0 | | | 0 |
| Mismatch rate per base, % | 0.00% | 0.11% | | 0.14% | | | 0.17% |
| Deletion rate per base | 0.01% | 0.01% | | 0.01% | | | 0.01% |
| Deletion average length | 1.853 | 1.8765 | | 1.8905 | | | 1.9015 |
| Insertion rate per base | 0.00% | 0.00% | | 0.00% | | | 0.00% |
| Insertion average length | 1.3475 | 1.293 | | 1.301 | | | 1.3095 |
| Number of reads mapped to multiple loci | 3,271,287.2 (2.64%) | 3,349,456.65 (2.70%) | | 3,367,742.45 (2.72%) | | | 3,375,106.85 (2.72%) |
| Number of reads mapped to too many loci | 534,171.1 (0.46%) | 503,661 (0.43%) | | 505,462.35 (0.43%) | | | 505,385.35 (0.43%) |
| Number of reads unmapped: too many mismatches | 0 (0.00%) | 0 (0.00%) | | 0 (0.00%) | | | 0 (0.00%) |
| Number of reads unmapped: too short | 8,046,584.8 (6.90%) | 5,332,962.25 (4.55%) | | 4,600,120.85 (3.91%) | | | 4,209,269.9 (3.57%) |
| Number of reads unmapped: other | 155,314.15 (0.12%) | 155,314.15 (0.12%) | | 155,314.15 (0.12%) | | | 155,314.15 (0.12%) |
| Number of chimeric reads | 0 (0.00%) | 0 (0.00%) | | 0 (0.00%) | | | 0 (0.00%) |
| **Morex-ref mapping (allowed max mismatch)** | **Morex (m=0)** | **Morex (m=2)** | | **Morex (m=4)** | | | **Morex (m=6)** |
| Number of input reads | 119,777,808.4 | 119,777,808.4 | | 119,777,808.4 | | | 119,777,808.4 |
| Average input read length | 298.25 | 298.25 | | 298.25 | | | 298.25 |
| Uniquely mapped reads number | 99,142,457.6 (82.63%) | 107,619,513 (89.78%) | | 109,509,789.6 (91.39%) | | | 110,294,053.35 (92.07%) |
| Average mapped length | 279.57 | 291.3445 | | 293.8135 | | | 294.656 |
| Number of splices: Total | 85,817,238.65 | 94,715,974.05 | | 96,584,061.6 | | | 97,226,134.45 |
| Number of splices: GT/AG | 84,565,642.5 | 93,343,681.8 | | 95,185,772.25 | | | 95,816,935.15 |
| Number of splices: GC/AG | 1,184,021.35 | 1,300,169.55 | | 1,325,371.35 | | | 1,335,853.25 |
| Number of splices: AT/AC | 67,574.8 | 72,122.7 | | 72,918 | | | 73,346.05 |
| Number of splices: Non-canonical | 0 | 0 | | 0 | | | 0 |
| Mismatch rate per base, % | 0.00% | 0.22% | | 0.30% | | | 0.35% |
| Deletion rate per base | 0.02% | 0.03% | | 0.03% | | | 0.03% |
| Deletion average length | 3.123 | 3.355 | | 3.453 | | | 3.49 |
| Insertion rate per base | 0.02% | 0.02% | | 0.02% | | | 0.02% |
| Insertion average length | 2.667 | 2.896 | | 2.9755 | | | 3.0205 |
| Number of reads mapped to multiple loci | 3,656,786.55 (3.01%) | 4,158,723.5 (3.43%) | | 4,232,954.1 (3.49%) | | | 42,751,39.5 (3.52%) |
| Number of reads mapped to too many loci | 131,798.65 (0.12%) | 103,843.4 (0.09%) | | 103,854.4 (0.09%) | | | 103,847.15 (0.09%) |
| Number of reads unmapped: too many mismatches | 0 (0.00%) | 0 (0.00%) | | 0 (0.00%) | | | 0 (0.00%) |
| Number of reads unmapped: too short | 16,700,021.9 (14.13%) | 7,748,984.8 (6.59%) | | 5,784,466.6 (4.91%) | | | 4,958,024.7 (4.20%) |
| Number of reads unmapped: other | 146,743.7 (0.12%) | 146,743.7 (0.12%) | | 146,743.7 (0.12%) | | | 146,743.7 (0.12%) |
| Number of chimeric reads | 0 (0.00%) | 0 (0.00%) | | 0 (0.00%) | | | 0 (0.00%) |

Table S4: Basic statistics of the sRTD and cRTD assembled from read alignment by allowing different mismatches.

| **Stat** | **sRTD (m=0)** | **sRTD (m=2)** | **sRTD (m=4)** | **sRTD (m=6)** | **cRTD (m=0)** | **cRTD (m=2)** | **cRTD (m=4)** | **cRTD (m=6)** |
| --- | --- | --- | --- | --- | --- | --- | --- | --- |
| Genome covered bases | 103,793,515 | 105,023,480 | 105,246,848 | 105,511,736 | 91,164,519 | 98,578,708 | 101,145,782 | 102,475,749 |
| Gene number | 59,447 | 60,787 | 61,246 | 61,469 | 59,664 | 64,176 | 66,741 | 68,225 |
| Multi-isoform gene number | 16,163 (27.19%) | 16,143 (26.56%) | 16,111 (26.31%) | 16,152 (26.28%) | 14,871 (24.92%) | 14,610 (22.77%) | 14,496 (21.72%) | 14,479 (21.22%) |
| Mono-exon gene number | 32,833 (55.23%) | 34,143 (56.17%) | 34,585 (56.47%) | 34,794 (56.6%) | 33,001 (55.31%) | 38,430 (59.88%) | 41,174 (61.69%) | 42,724 (62.62%) |
| Multi-exon gene number | 26,614 (44.77%) | 26,644 (43.83%) | 26,661 (43.53%) | 26,675 (43.4%) | 26,663 (44.69%) | 25,746 (40.12%) | 25,567 (38.31%) | 25,501 (37.38%) |
| ^a^ Protein-coding | 33,254 (55.94%) | 33,394 (54.94%) | 33,473 (54.65%) | 33,496 (54.49%) | 31,961 (53.57%) | 32,570 (50.75%) | 32,959 (49.38%) | 33,156 (48.6%) |
| ^b^ Best-hit in UniProt Plant (e < 0.01) | 21,365 (35.94%) | 21,425 (35.25%) | 21,435 (35%) | 21,455 (34.9%) | 20,893 (35.02%) | 20,821 (32.44%) | 20,824 (31.2%) | 20,876 (30.6%) |
| Transcript number | 144,872 | 145,906 | 146,448 | 146,727 | 128,438 | 133,595 | 135,772 | 137,353 |
| Mono-exon transcript number | 33,429 (23.07%) | 34,743 (23.81%) | 35,163 (24.01%) | 35,374 (24.11%) | 33,578 (26.14%) | 38,973 (29.17%) | 41,707 (30.72%) | 43,276 (31.51%) |
| Multi-exon transcript number | 111,443 (76.93%) | 111,163 (76.19%) | 111,285 (75.99%) | 111,353 (75.89%) | 94,860 (73.86%) | 94,622 (70.83%) | 94,065 (69.28%) | 94,077 (68.49%) |
| Protein-coding | 73,998 (51.08%) | 74,227 (50.87%) | 74,553 (50.91%) | 74,415 (50.72%) | 64,294 (50.06%) | 65,513 (49.04%) | 65,770 (48.44%) | 65,952 (48.02%) |
| Best-hit in UniProt Plant (e < 0.01) | 47,614 (32.87%) | 47,699 (32.69%) | 47,836 (32.66%) | 47,763 (32.55%) | 42,126 (32.8%) | 42,227 (31.61%) | 42,036 (30.96%) | 42,057 (30.62%) |
| Protein average length | 399.93 | 399.57 | 398.62 | 398.52 | 383.5 | 384.56 | 383.5 | 382.98 |
| Transcript number per gene | 2.44 | 2.4 | 2.39 | 2.39 | 2.15 | 2.08 | 2.03 | 2.01 |
| Transcript N50 | 3,284 | 3,303 | 3,293 | 3,295 | 3,056 | 3,185 | 3,191 | 3,192 |
| Transcript N90 | 1,457 | 1,452 | 1,450 | 1,450 | 1,286 | 1,330 | 1,325 | 1,317 |
| Transcript average length (exonic) | 2486.78 | 2484.39 | 2473.73 | 2473.15 | 2218.22 | 2275.02 | 2257.67 | 2244.46 |
| Transcript total length (exonic) | 360,264,709 | 362,486,762 | 362,272,529 | 362,878,452 | 284,903,601 | 303,931,318 | 306,528,756 | 308,283,084 |
| Exon number | 881,447 | 876,865 | 878,078 | 877,283 | 722,816 | 726,643 | 724,892 | 726,459 |
| Exon number per transcript | 6.08 | 6.01 | 6 | 5.98 | 5.63 | 5.44 | 5.34 | 5.29 |
| Exon average length | 408.72 | 413.39 | 412.57 | 413.64 | 394.16 | 418.27 | 422.86 | 424.36 |
| Intron number | 736,575 | 730,959 | 731,630 | 730,556 | 594,378 | 593,048 | 589,120 | 589,106 |
| Intron number per transcript | 5.08 | 5.01 | 5 | 4.98 | 4.63 | 4.44 | 4.34 | 4.29 |
| Intron average length | 643.87 | 648.8 | 648.38 | 648.35 | 655.84 | 659.47 | 660.57 | 659.86 |

Table S5: The misassembled transcripts filtered at the quality control step.

| **Transcript category** | **sRTD (m=0)** | **sRTD (m=2)** | **sRTD (m=4)** | **sRTD (m=6)** | **cRTD (m=0)** | **cRTD (m=2)** | **cRTD (m=4)** | **cRTD (m=6)** |
| --- | --- | --- | --- | --- | --- | --- | --- | --- |
| Unstranded | 106 | 135 | 126 | 131 | 123 | 136 | 144 | 147 |
| Non canonical SJ | **106** | **179** | **302** | **350** | **86** | **2857** | **4041** | **4454** |
| Low read support SJ | 16436 | 16456 | 16509 | 16524 | 18407 | 20543 | 20747 | 20837 |
| Redundant | 1 | 1 | 1 | 1 | 0 | 0 | 0 | 0 |
| Fragment | 14063 | 14217 | 14124 | 14053 | 12521 | 11881 | 11903 | 11770 |

Table S6: Gffcompare structure comparisons of sRTD with 0 mismatch and cRTD with 2 mismatches.

| **Genomic regions** | | | cRTD.minimap2_m0 vs  sRTD_m0 | | | cRTD.minimap2_m2 vs  sRTD_m0 | |
| --- | --- | --- | --- | --- | --- | --- | --- |
| False negative | Missed exons | | 31391/291963 (10.8%) | | | 32335/291963 (11.1%) | |
|  | Missed introns | | 18213/172487 (10.6%) | | | 21024/172487 (12.2%) | |
|  | Missed loci | | 12787/59440 (21.5%) | | | 10808/59440 (18.2%) | |
| False positive | Novel exons | | 12815/259285 (4.9%) | | | 16897/264186 (6.4%) | |
|  | Novel introns | | 4360/154670 (2.8%) | | | 4853/152644 (3.2%) | |
|  | Novel loci | | 7848/57517 (13.6%) | | | 11448/61568 (18.6%) | |
| **Individual transcripts** | | cRTD.minimap2_m0 vs sRTD_m0 | | | cRTD.minimap2_m2 vs sRTD_m0 | | |
| Statistics | | Recall | | Precision | Recall | | Precision |
| Base level | | 76.9% | | 90.2% | 80.1% | | 87.4% |
| Exon level | | 74.1% | | 81.5% | 74.9% | | 81.4% |
| Intron level | | 84.6% | | 94.4% | 82.8% | | 93.5% |
| Intron chain level | | 57.6% | | 68.6% | 53.6% | | 64.2% |
| Transcript level | | 56.8% | | 65.1% | 55.8% | | 61.6% |
| Locus level | | 62.5% | | 64.6% | 67.3% | | 65.0% |

Summary for gene loci (clusters of exon overlaps):

a) Labels m0 and m2: assembled RTDs with 0 and 2 mismatches in the read mapping.
b) The assembled cRTD with 0 and 2 mismatches were aligned to Barke genome with minimpa2, respectively.
c) Reference sRTD: 144872 transcripts in 59440 loci (111443 multi-exon)
d) Query cRTD.minimap2_m0: 126532 transcripts in 57517 loci (93268 multi-exon transcripts)
e) Query cRTD.minimap2_m2: 131223 transcripts in 61568 loci (93036 multi-exon transcripts)
